# Induced CD45 Proximity Potentiates Natural Killer Cell Receptor Antagonism

**DOI:** 10.1101/2022.06.27.497842

**Authors:** Junming Ren, Yeara Jo, Lora K. Picton, Leon L. Su, David H Raulet, K. Christopher Garcia

**Affiliations:** Department of Molecular and Cellular Physiology, Stanford University School of Medicine, Stanford, United States; Howard Hughes Medical Institute, Stanford University School of Medicine, Stanford, United States; Division of Immunology and Molecular Medicine, Department of Molecular and Cell Biology, University of California, Berkeley, Berkeley, United States

**Keywords:** natural killer cell receptors, CD45, bispecific antibodies, protein engineering

## Abstract

Natural Killer (NK) cells are a major subset of innate immune cells that are essential for host defense against pathogens and cancer. Two main classes of inhibitory NK receptors (NKR), KIR and CD94/NKG2A, play a key role in suppressing NK activity upon engagement with tumor cells or virus infected cells, limiting their antitumor and antiviral activity. Here, we find that single-chain mouse NKR antagonists linked to a VHH that binds the cell surface phosphatase CD45 potentiate NK and T activity to a greater extent than NKR blocking antibodies alone *in vitro*. We also uncovered crosstalk between mouse NKG2A and Ly49 that collectively inhibit NK cell activation, such that CD45-NKG2A and CD45-Ly49 bispecific molecules show synergistic effects in their ability to enhance NK cell activation. The basis of the activity enhancement by CD45 ligation may reflect greater antagonism of inhibitory signaling from engagement of MHC I on target cells, combined with other mechanisms, including avidity effects, tonic signaling, antagonism of weak inhibition from engagement of MHC I on non-target cells and possibly CD45 segregation within the NK cell-target cell synapse. These engineered ligands uncover a mechanism for enhancing the activity of mouse NK and T cells that merits evaluation in the context of human NKR antagonist cancer immunotherapies.

## Introduction

Natural Killer (NK) cells and T cells are lymphocytes that protect against infection and cancer (*1*, *2*). The function of NK cells is tightly regulated by two classes of NK receptors, inhibitory receptors and activating NK receptors, which engage ligands such as MHC I and in turn tune the level of NK cell activity. Dysregulation of the balance between inhibitory and activating signaling in the context of tumor microenvironments or viral infection results in reduced NK effector function (*3*–*9*). NKG2A and the family of Ly49 receptors are the two major inhibitory NK receptor types in mice, which have inhibitory tyrosine-based inhibition motifs (ITIMs) present in their intracellular domains (ICD) (*4*, *10*–*13*). One or more NKG2A and/or Ly49 family receptors are expressed on a large percentage of NK cells, as well as a small subset of CD8^+^T cells (*14*–*20*). Most activated CD8 T cells upregulate NKG2A but not Ly49 recepters (*20*). NKG2A forms a heterodimer with CD94, whereas Ly49 receptors are displayed as homodimers on the surface of lymphocytes. Upregulation of inhibitory NK receptors and their ligands is frequently observed in different diseases (*21*, *22*). Engagement of NKG2A/CD94 or Ly49C/I receptors with MHC I molecules induces the phosphorylation of ITIM motifs in NK receptors, resulting in phosphatase binding. Recruitment of Src homology 1-containing phosphotyrosine phosphatase (SHP1) results in the transmission of inhibitory signals that repress activating NK signaling (*23*). In humans, NKG2A and the counterpart of Ly49 receptors, Inhibitory killer cell immunoglobulin-like receptors (KIR), play a key role in suppressing NK and T cell activity by interacting with different Human Leukocyte Antigens (HLAs), including HLA-E, HLA-A, HLA-B and HLA-C (*24*).

Monoclonal antibody (mAb) based immune checkpoint blockade (ICB) has revolutionized the treatment of cancer. Checkpoint inhibitors result in durable antitumor effects in many patients with metastatic and treatment-refractory cancers (*25*–*28*). ITIM bearing receptors including PD-1, CTLA-4 and NKG2A have been identified as key IgSF molecules on both T cells and NK cells (*14*, *15*, *29*–*32*). Inhibition of mouse NKG2A with the 20D5 mAb restores the effector function of NK and T cells in lung cancer and lymphoma tumor models (*14*, *15*). Antibodies to inhibitory KIR are being evaluated for the treatment of cancer, and two mAbs are being tested in clinical trials in lymphoma and leukemia. IPH4102, specific for KIR3DL2, is in a clinical trial of T Cell Lymphoma, whereas Lirilumab, specific for KIR2DL1-3, has not shown good efficacy in its phase II trial for the treatment of acute myeloid leukemia (AML)(*33*, *34*). So far, anti-KIR mAbs have yet to show robust efficacy in clinical trials (*33*–*36*), suggesting that additional innovation may be required to target this interaction more effectively for cancer immunotherapy.

CD45 is a cell surface phosphatase ubiquitously expressed on immune cells, that serves to modulate the extent of phosphorylation of many immunoreceptors through a variety of mechanisms including direct dephosphorylation and segregation from the immune synapse (*37*, *38*). Despite its central role in the signaling response at the membrane of immune cells, CD45 has not been actively investigated as an immunotherapeutic target. In one approach, <u>R</u>eceptor <u>I</u>nhibition by <u>P</u>hosphatase <u>R</u>ecruitment (RIPR), cis-ligation of CD45 with the PD-1 checkpoint receptor using bispecific molecules potentiated inhibition of the checkpoint activity (*39*). By cross-linking PD-1 to CD45, the PD-1-RIPR suppressed both tonic and ligand-activated signaling in T cells to enhance T cell effector functions. Additional mechanisms may also play a role, such as “passenger-like” sequestration of the PD1-CD45 conjugate away from the immune synapse. Here we asked if CD45-ligation to inhibitory NKR could potentiate NKR antagonism, more effectively reversing inhibition through these receptors. We find that a series of CD45-targeted NK receptor antagonists potentiate mouse NK and CD8^+^ T cell activity *in vitro*. We also find synergy between Ly49 and NKG2A antagonism on mouse NK cells, suggesting that maximally efficacious NKR targeting may require combination approaches. This ligand engineering approach warrants consideration for enhanced human NK receptor antagonism.

## Results

Ligation of MHC I or Qa-1^b^ to Ly49 or NKG2A receptors on NK cells, respectively, results in strong inhibition of NK activities (*15*, *40*, *41*). The MHC molecules present in the C57BL/6 mice examined in this study engage Ly49C, Ly49I and NKG2A. To compare the impacts of blocking Ly49I or NKG2A on NK function, we sorted Ly49I^+^ or NKG2A^+^ NK cells and cocultured them with RMA tumor cells, which express MHC I. The RMA cells for coculturing with NKG2A^+^ NK cells were pre-cultured in IFNγ to induce Qa-1^b^ expression. Target cell lysis by Ly49I^+^ NK cells was modestly augmented by the Ly49C/I-specific 5E6-mAb, whereas lysis by NKG2A^+^ NK cells was modestly augmented by the NKG2A-C specific 20D5-mAb but not by another NKG2A-specific antibody, 16A11, which may not interfere with binding to Qa-1^b^ (Figure 1A, B, S1A). Both 5E6-mAb and 20D5-mAb bound well to primary NK cells (Figure 1C, D). Given that nearly all NK cells that express NKG2A or Ly49I also express CD45 (Figure 1E), we reengineered 20D5 and 5E6 antibodies into bispecific molecules by fusing an anti-CD45 single domain antibody fragment (VHH) to the N terminus of 20D5 or 5E6-derived scFvs using a flexible (Gly-Ser)_4_ linker. (Fig 1F, I). We sequenced the VH and VL of antibodies from hybridomas producing these antibodies, and re-formatted VH and VL domains into scFv, as well as CD45 VHH-NKR scFv fused molecules (Figure S1B). Following the production of these recombinant proteins from insect cells (Fig 1G, H, J, K), we checked the cell binding of 5E6-scFv and 20D5-scFv in Ly49^+^ and NKG2A^+^ NK cells, and observed dose-dependent binding to NK cells, with 20D5-scFv appearing to be higher affinity (Fig 1L).

**Figure 1.**
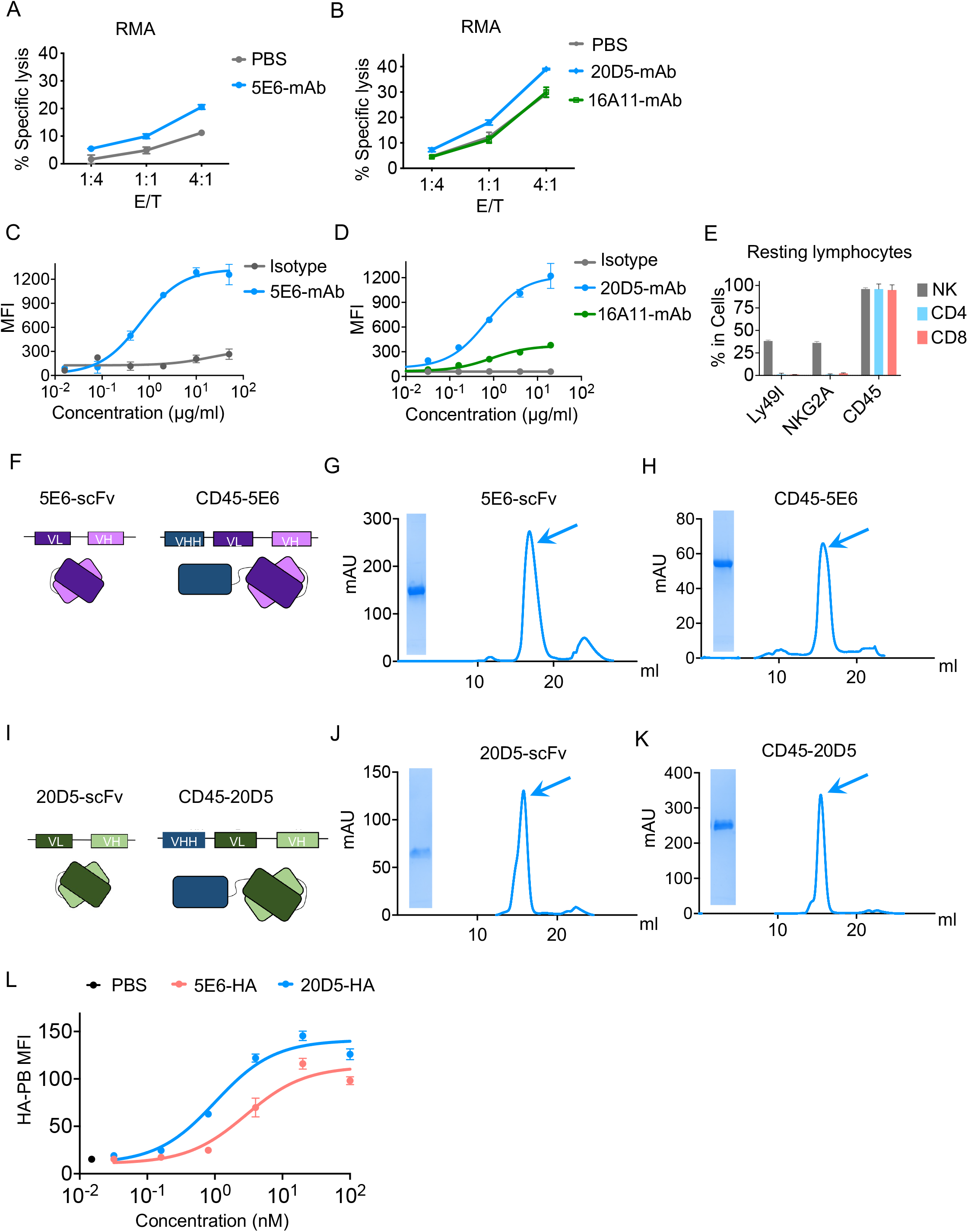
Engineering strategy of CD45-NKR. (A) Cytolysis of RMA cells expressing MHC I by sorted Ly49I^+^ NK cells in the presence or absence of 100nM Ly49C/I specific 5E6-mAb were tested by Annexin V/7-AAD staining. NK cells were expanded by 100IU mouse IL-2. Ly49I^+^ NK cells were sorted and allowed to recover for 2 days. (B) NK cells were tested for cytolysis of RMA cells in the presence or absence of NKG2A specific 16A11-mAb and 20D5-mAb. NK cells were expanded by 100IU mouse IL-2 for 5 days. NKG2A^+^ NK cells were sorted and allowed to recover for 2 days. RMA cells were stimulated with 10ng/μl IFNγ for 2 days for upregulating Qa-1^b^. (C) Representative titration of 5E6-mAb on sorted mouse NK cells. The titration data shown are median fluorescence intensity (MFI) over a range of anti IgG-AF647 antibody concentrations. (D) Representative titration of 20D5-mAb vs 16A11 mAb on sorted mouse NK cells. The titration data shown are MFI over a range of anti IgG-AF647 antibody concentrations. (E) Splenocytes from wild-type C57BL/6 mice were harvested and stained with a panel of antibodies against lymphocyte markers, the cells were analyzed by flow cytometry. Quantification of percentage of Ly49I, NKG2A and CD45 expression on splenic NK cells (CD45.2^+^, CD3^-^, NK1.1^+^), CD4^+^ T cells (NK1.1^-^, CD3^+^, CD4^+^) and CD8^+^ T cells (NK1.1^-^, CD3^+^, CD8^+^). (F) Schematic representation of 5E6-scFv and CD45-5E6 depicting the connection between the CD45 VHH and variable heavy (VH) and variable light (VL) chains of 5E6-scFv. (G, H) Size exclusion chromatography profiles of 5E6-scFv (G) and CD45-5E6 (H). Arrowheads indicate the collected fraction analyzed by SDS-PAGE. (I) Schematic representation of 20D5-scFv and CD45-20D5 depicting the connection between the CD45 VHH and variable heavy (VH) and variable light (VL) chains of 20D5-scFv. (J, K) SEC profiles of 20D5-scFv (J) and CD45-20D5 (K). Arrowheads indicate the collected fraction analyzed by SDS-PAGE. (L) Cell-surface binding of 5E6-scFv or 20D5-scFv. Mouse primary NK cells were incubated with varying concentration of HA-tagged 5E6-scFv or 20D5-scFv, washed, and stained with anti-HA-PB, then analyzed via flow cytometry. All curves show mean ± Standard deviation (SD) of n=3 per group and data are representative of two independent experiments.

We tested whether the Ly49-targeted CD45-5E6 could enhance NK-mediated cell killing (Figure 2A, B). Notably, when sorted Ly49I^+^ NK cells were cocultured with MHC I^+^ RMA cells, the Ly49C/I blocking 5E6-scFv augmented target cell killing by 7% at the highest concentration tested, whereas the Ly49C/I-targeting CD45-5E6 augmented target cell killing by more than 20% (Figure 2A, B, S2A, B). We next determined the effects of CD45-5E6 on NK activation and degranulation. Recruitment of SHP1 by the MHC I-Ly49 signaling axis results in suppression of proximal NK activating receptor signaling, which is indicated by CD69 upregulation, and downstream effects, including degranulation and cytokine production. In a NK and RMA target cell coculture assay, we performed a dose-series stimulation of CD45-5E6 and 5E6-scFv molecules. Compared to 5E6-scFv, CD45-5E6 was much more efficient at augmenting degranulation and IFNg production (the latter determined by intracellular cytokine staining), as well as induction of CD69 expression, by Ly49I^+^ NK cells (Figure 2C-E, S2C). To summarize, we re-engineered the antibodies against NKG2A or Ly49C/I into CD45-targeted versions, which show a superior capacity to enhance NK cytolysis, degranulation, IFNγ production and NK activating receptor signaling compared to NKR blocking antibodies alone. From these experiments, it is not clear whether the effect is due to more efficient interference with inhibition mediated by interactions with target cell MHC I, repression of tonic signaling, avidity-based due to two-site binding, or possibly enhanced segregation of NKR from the NK cell synapse.

**Figure 2.**
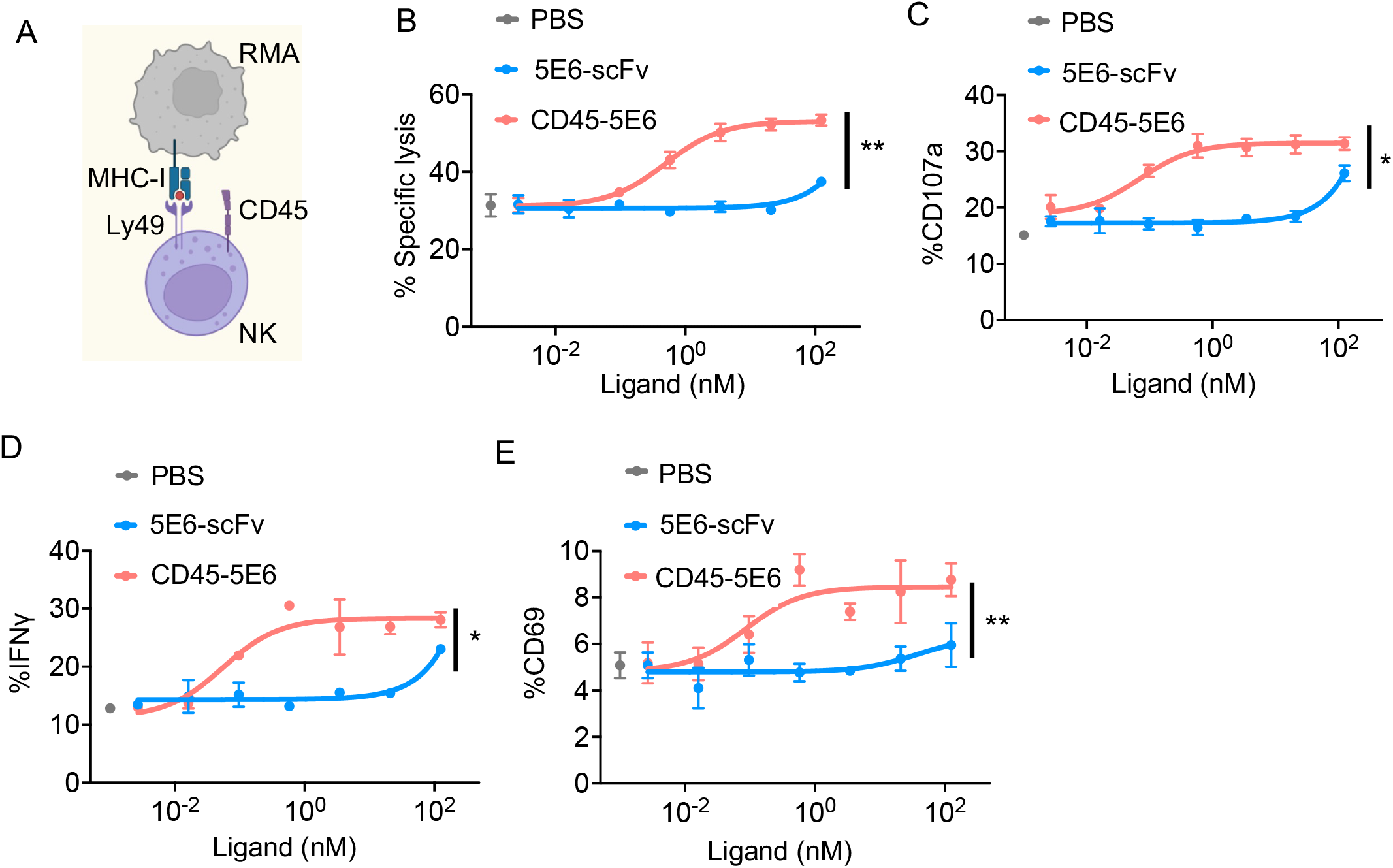
Functional properties of CD45-5E6 in mouse NK cells. (A) Schematics of NK cell killing assay of RMA cells expressing MHC I. (B) Sorted Ly49I^+^ NK cells were co-cultured with RMA cells in the presence or absence of 5E6-scFv or CD45-5E6 for 4h and assayed for dose dependent cytolysis of RMA cells by Annexin V/7-AAD staining. (C) IL-2 expanded NK cells were co-cultured with RMA cells in the presence or absence of 5E6-scFv or CD45-5E6 for 4h and assayed for dose dependent degranulation by cell surface staining of CD107a in Ly49I^+^ NK cells. (D) IL-2 expanded NK cells were co-cultured with RMA cells in the presence or absence of 5E6-scFv or CD45-5E6 for 4h and assayed for dose dependent IFNγ secretion by intracellular staining of IFNγ in Ly49I^+^ NK cells. (E) IL-2 expanded NK cells were co-cultured with RMA cells in the presence or absence of 5E6-scFv or CD45-5E6 for 4h and assayed for dose dependent degranulation by cell surface staining of CD69 in Ly49I^+^ NK cells. All curves show mean ± SD of n=3 per group. Groups were compared by unpaired t-test and data are representative of two independent experiments. *P < 0.05; **P < 0.01; ***P < 0.001; ****P < 0.0001.

NKG2A is expressed by primary resting and IL-2 cultured NK cells to a similar extent as Ly49I (Figure 3A). We therefore employed the same strategy as used to generate the CD45-5E6 bispecific molecule to generate and test an NKG2A-targeted CD45-20D5 bispecific molecule (Figure 1I-K, Figure 3B). Notably, treatment with CD45-20D5 resulted in a higher maximal degranulation of NK cells compared to 20D5-scFv (Figure 3C). NKG2A is expressed by naïve NK cells and not naïve CD8 T cells, but is strongly upregulated in virus-specific CD8^+^T cells and tumor antigen specific T cells(*15*, *20*). To examine whether CD45-20D5 plays a role in CD8^+^ T cells, we tested its effects in a system of MHC class I-restricted, ovalbumin-specific, TCR transgenic CD8^+^ T cells (OT-I) and B16-OVA target cells, which had been cultured with IFNγ to induce MHC I and Qa-1^b^ (Fig. S3A-E). In this system, the mouse B16F10 melanoma cells express a model antigen, chicken ovalbumin, and therefore present the OVA peptide antigen and activate OVA specific CD8^+^ OT-I cells. For generating effector OT-I cells, splenocytes from OT-I mice were stimulated with anti CD28 and OVA_257-264_ octapeptide, SIINFEKL, for 8 days. Although NKG2A is expressed at low levels in resting OT-I cells, it was robustly upregulated in effector OT-I cells following activation (Figure 3E). Surface staining of CD107a, CD25 and CD69 on OT-I cells following coculture with B16-OVA cells indicated that CD45-20D5 promotes greater OT-I degranulation and induces greater TCR signaling than 20D5-scFv (Figure 3F-H, S3F). Moreover, CD45-20D5 stimulation resulted in greater production of the effector cytokines IL-2 and IFNγ as compared to 20D5-scFv (Figure 3I-J).

**Figure 3.**
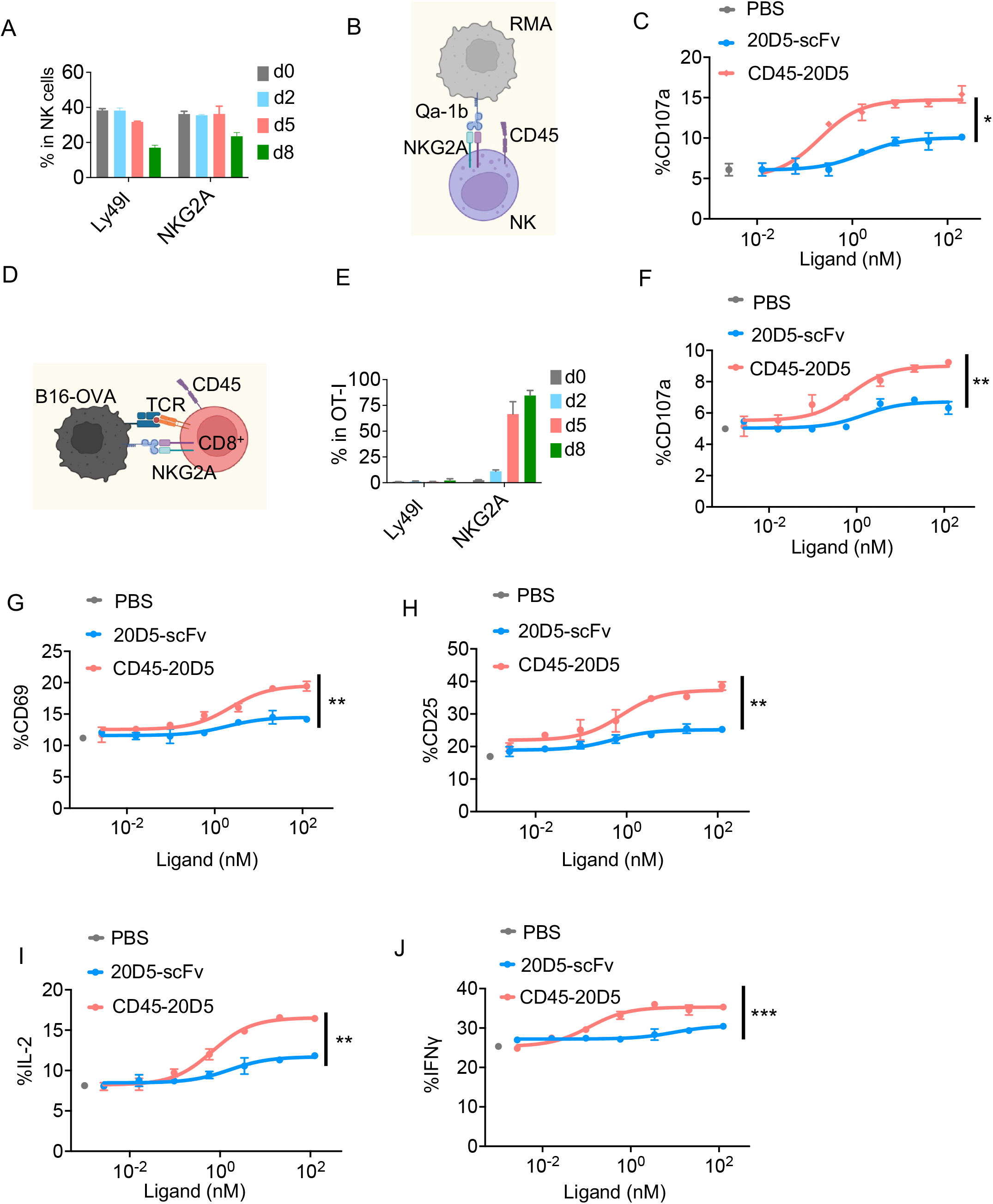
Functional properties of CD45-20D5 in mouse NK cells and CD8^+^ T cells. (A) Quantification of percentage of Ly49I^+^ and NKG2A^+^ NK cells cultured in 100IU mouse IL-2 for days indicated. (B) Schematic of co-culture of NKG2A^+^ NK cells with RMA cells expressing Qa-1^b^. (C) RMA cells were treated with IFNγ for upregulating Qa-1^b^. IL-2 expanded NK cells were cocultured with RMA cells in the presence or absence of 20D5-scFv or CD45-20D5 for 4h and assayed for dose dependent degranulation by cell surface staining of CD107a in NKG2A+ NK cells. (D) Schematic of co-culture of NKG2A^+^ OT-I cells with B16-OVA cells expressing Qa-1^b^. (E) Quantification of percentage of Ly49I and NKG2A expression during generating OT-I effector cells by incubating with 0.5μg/ml anti-CD28 and 1μg/ml SIINFEKL peptides. (F) B16-OVA cells were treated with IFNγ for upregulating Qa-1^b^. Purified CD8^+^ OT-I cells were co-cultured with B16-OVA cells in the presence or absence of 20D5-scFv and CD45-20D5 for 4h and assayed for dose dependent degranulation by cell surface staining of CD107a in NKG2A^+^ OT-I cells. (G, H) Dose-response curves showing CD69 (G) and CD25 (H) expression in response to20D5-scFv and CD45-20D5 stimulation. (I, J) Dose-response curves showing intracellular staining of IL-2 (I) and IFNγ (J) in response to 20D5-scFv and CD45-20D5 stimulation. All curves show mean ± SD of n=3 per group. Groups were compared by unpaired t-test and data are representative of two independent experiments. *P < 0.05; **P < 0.01; ***P < 0.001; ****P < 0.0001.

Previous studies have shown that many T, B, and NK cell receptors can in some cases signal even when the ligand is absent on the target cell (*32*, *35*–*37*). Such effects may have different explanations, including tonic signaling, where the receptor signals independently of ligand, cis-signaling, where the receptor engages the ligand on the membrane of the NK or T cell ltself, or possibly when the receptor is engaged by ligand on a third party cell in the vicinity. To investigate whether the impact of CD45-NKR is fully dependent on target cell engagement of the NK receptor ligand we first examined the expression of Qa-1^b^ and MHC I on NK cells by flow cytometry and found that both Qa-1^b^ and MHC I can be detected on the surface of NK cells (Fig. S4A, B). We used a peptide-MHC I deficient target cell line, RMA-*B2m* KO, to address whether the impact of CD45-NKRs is dependent on MHC I expression by target cells (Figure 4A, S4C, D). We found that the CD45-5E6 significantly augmented killing of MHC I deficient RMA cells (Fig. 4A) albeit to a smaller extent than it augmented killing of MHC I^+^ RMA cells (Fig. 4B). 5E6-scFv had no effect. Similarly, CD45-20D5 modestly augmented the killing of RMA target cells that lacked Qa-1^b^ expression (RMA-Qa-1^b^ KO) (Fig. 4C) albeit to a smaller extent that it augmented killing of RMA or B16 target cells that expressed Qa-1^b^ (Fig. 4D, E). 20D5-scFv, in contrast, did not augment killing of Qa-1^b^-deficient cells but did augment killing of WT cells. These data suggest that enhanced antagonism by CD45-NKRs reflects antagonism of at least two separable inhibitory effects of the NK receptors: inhibitory signaling resulting from engagement of MHC I on target cells and inhibitory signaling that is independent of target cell MHC I expression. The weaker inhibition that is independent of target cell MHC I expression may be due to engagement of MHC I on other cells or could reflect “tonic” inhibitory signaling that the receptor confers without engaging MHC I.

**Figure 4.**
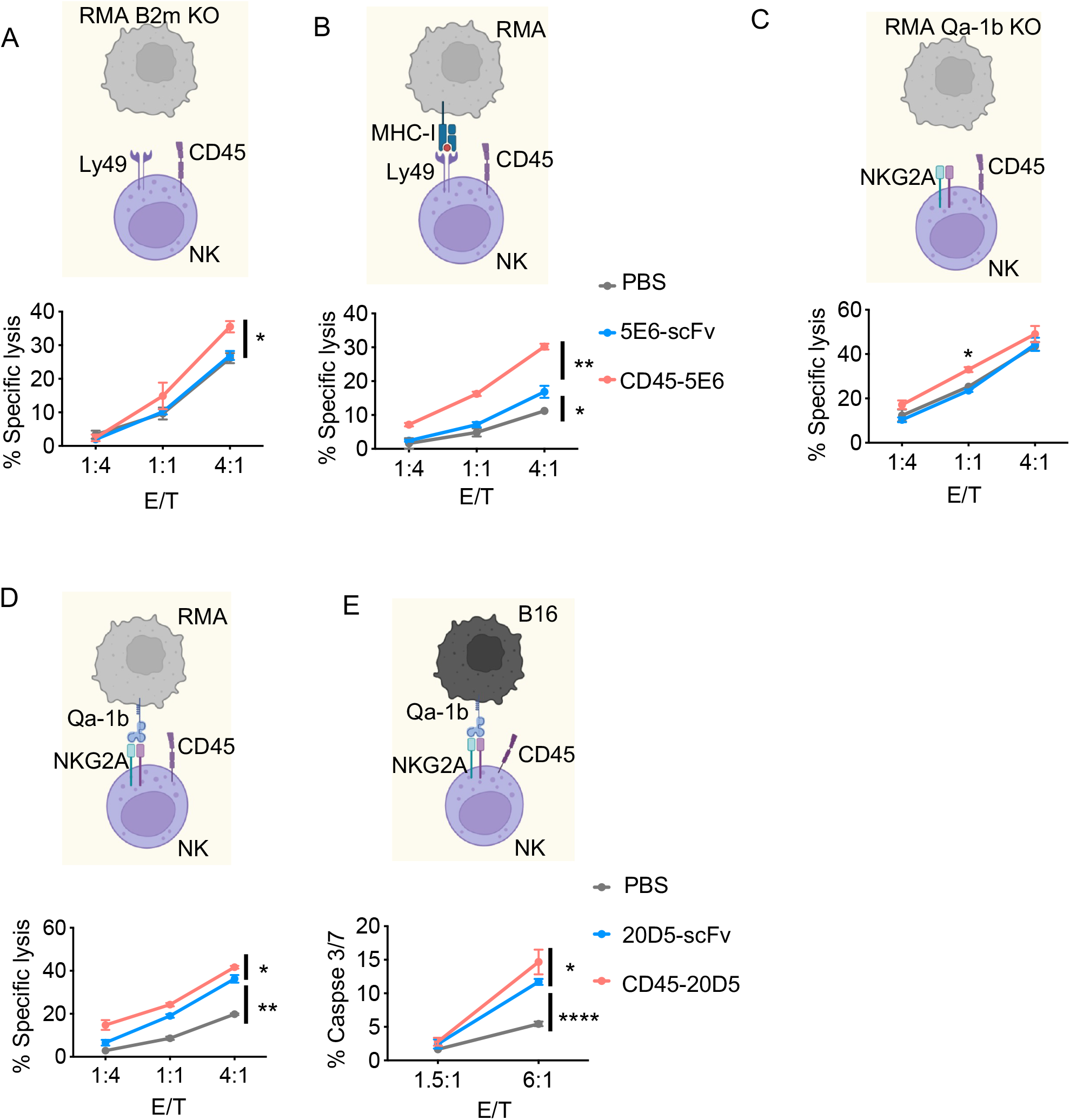
CD45-NKR potentiates NK killing of RMA cells. (A, B) NK cells were expanded by mouse IL-2 for 5 days. Ly49I^+^ NK cells were sorted and allowed to recover for 2 days. Cytolysis of RMA B2m KO cells (A), wildtype RMA cells (B) by sorted Ly49I^+^ NK cells in the presence or absence of 100nM 5E6-scFv or CD45-5E6. Percent lysis determined by Annexin V/7-AAD staining. (C, D) NKG2A^+^ NK cells were sorted and allowed to recover for 2 days. Wildtype RMA cells were treated with IFNγ for Qa-1^b^ upregulation. Cytolysis of RMA Qa-1^b^ KO (C), wildtype RMA cells (D) by NKG2A^+^ NK cells in the presence or absence of 20D5-scFv or CD45-20D5 at 100nM. (E) B16 cells were treated with IFNγ for Qa-1^b^ upregulation. Cytolysis of B16 cells by NKG2A^+^NK cells in the presence or absence of 20D5-scFv or CD45-20D5 at 100nM were tested by detecting Caspase-3/7 activity. All curves show mean ± SD (n=3) and were analyzed by one-way ANOVA relative to 5E6-scFv or 20D5-scFv. Multiple comparisons were corrected using Dunnett’s test. Data are representative of two independent experiments. *P < 0.05; **P < 0.01; ***P < 0.001; ****P < 0.0001.

The activation state of NK cells is tightly regulated by the receptors they express, such that NK cells lacking all inhibitory receptors for self MHC I exist in a hyporesponsive state (*42*, *43*).Thus, we further investigated the impact of CD45-NKRs on responses of freshly isolated NK cells in cocultures with target cells. Preactivated NK cells were harvested from mice pretreated with Poly(I:C), and were cocultured with target cells (Fig. 5A, S5A). When compared to 5E6-scFv, CD45-5E6 induced a substantially greater response in Ly49I^+^ NK cells (Fig. 5B) Importantly, Ly49I^+^, but not Ly49I^-^, NK cells showed significantly enhanced responsiveness in the presence of CD45-5E6 (Fig. 5B, C). Moreover, CD45-5E6 stimulated many more of the Ly49I^+^ NK cells to produce effector cytokines and chemokines, such as IFNγ and CCL5, than did 5E6-scFv (Figure 5D, E).

**Figure 5.**
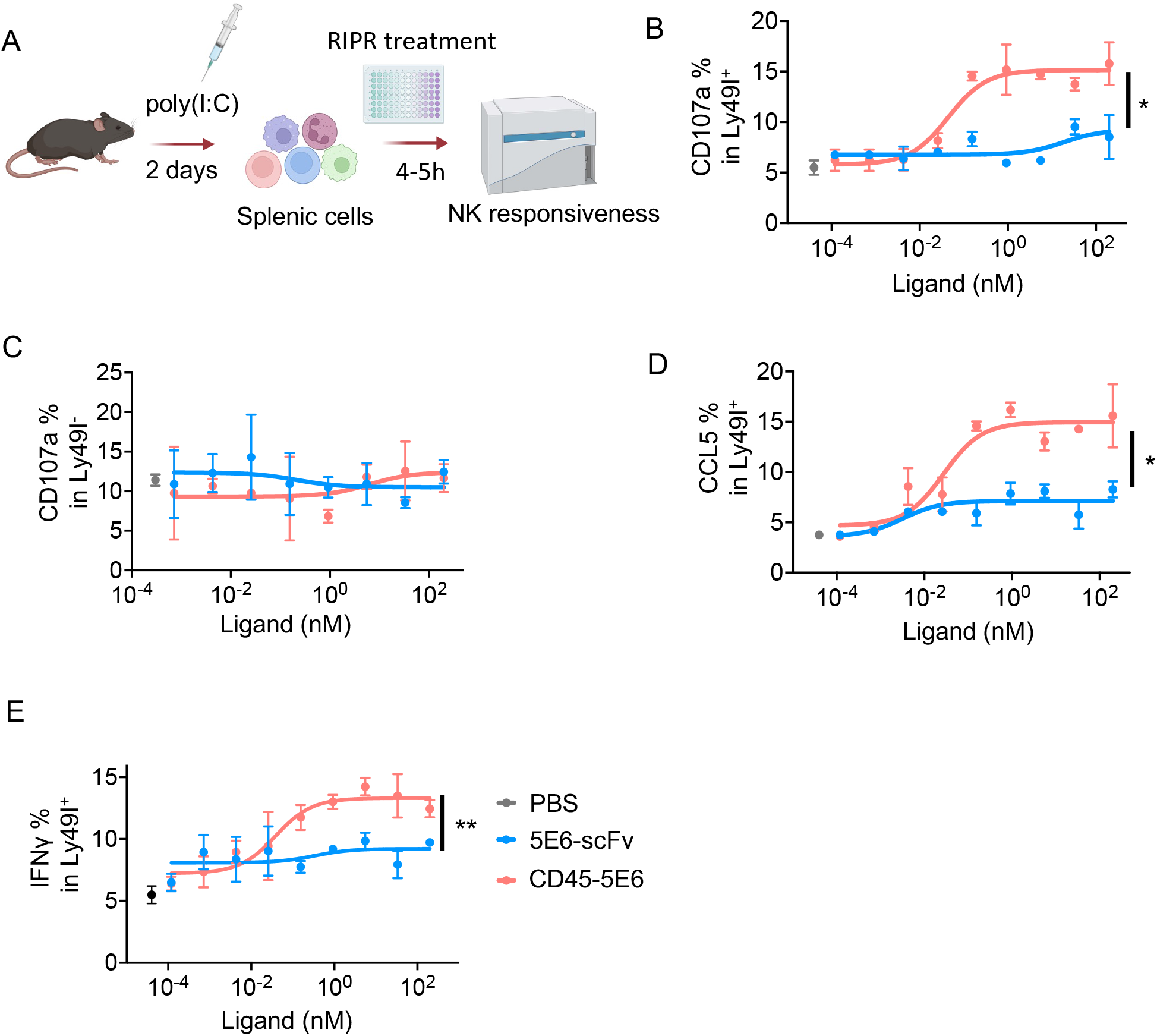
NKG2A suppresses Ly49C/I, combination of CD45-5E6 and CD45-20D5 synergizes NK activation. (A) Schematics of NK responsiveness assay. WT mice were treated with 200μg poly(I:C) for 2 days, splenocytes were harvested and cocultured with RMA cells in presence of 5E6-scFv or CD45-5E6 for 5h, NK responsiveness was tested by detecting surface expression of CD107a in Ly49I^+^ NK cells. (B, C) The NK cells were assayed for dose dependent CD107a in Ly49I^+^ (B) and Ly49I^-^ NK (C) cells by surface staining. (D, E) Dose-response curves showing percentage of CCL5^+^ (D) and CD107a^+^IFNγ^+^ (E) in Ly49I^+^ NK cells in response to 5E6-scFv or CD45-5E6 stimulation. Curves show mean ± SD (n=3) and were analyzed by one-way ANOVA relative to 5E6-scFv. Multiple comparisons were corrected using Dunnett’s test. Bar graphs show mean ± SD (n=4) and were analyzed by two-way ANOVA. Multiple comparisons were corrected using Tukey test. Data are representative of three independent experiments. *P < 0.05; **P < 0.01; ***P < 0.001; ****P < 0.0001.

Primary NK cells can be divided into eight distinct subpopulations based on the surface expression and co-expression of Ly49C, Ly49I and NKG2A (Figure 6A, B Fig. S6A). To address how these different NK cell subpopulations respond to the CD45-NKRs, we gated the cells stained with antibodies against Ly49C, Ly49I and NKG2A. The responses by each of the eight subpopulations of primary NK cells to CD45-NKRs and scFvs is shown in Fig. S6. Figure 6 shows a simpler depiction, representing the responses by four populations of NK cells (Fig. 6B): those that co-expressed NKG2A with either Ly49I and/or Ly49C; those that expressed Ly49I and/or Ly49C but not NKG2A; those that expressed NKG2A alone (Ly49C/I-negative); and those that lacked all three of these receptors. Interestingly, NK cells that co-expressed NKG2A with either Ly49I and/or Ly49C did not respond to either CD45-5E6 or CD45-20D5 separately but responded well when both CD45-NKRs were present (Fig. 6C, Fig. S6B, D, E). Treatments with CD45-5E6 alone potentiated responses of NK cells that expressed Ly49C and/or Ly49I, but only if they lacked NKG2A (Fig. 6D vs 6C, Fig. S6C, F, G). Conversely, treatments with CD45-20D5 alone potentiated responses of NK cells that expressed NKG2A, but only if they lacked Ly49C and Ly49I (Fig. 6E vs 6C, Fig. S6H). In contrast to the results with CD45-NKRs, 5E6-scFv was ineffective with Ly49C/I^+^NKG2A^-^ NK cells (Fig. 6H, Fig. S6C, F, G) and the combination of 5E6-scFv and 20D5-scFv was ineffective with Ly49C/I^+^NKG2A^+^NK cells (Fig. 6G, Fig. S6B, D, E). However, the 20D5 scFv had similar activity to CD45-20D5 in activating Ly49C/I^-^NKG2A^+^ NK cells (Fig. 6I vs 6E, Fig. S6G).

**Figure 6.**
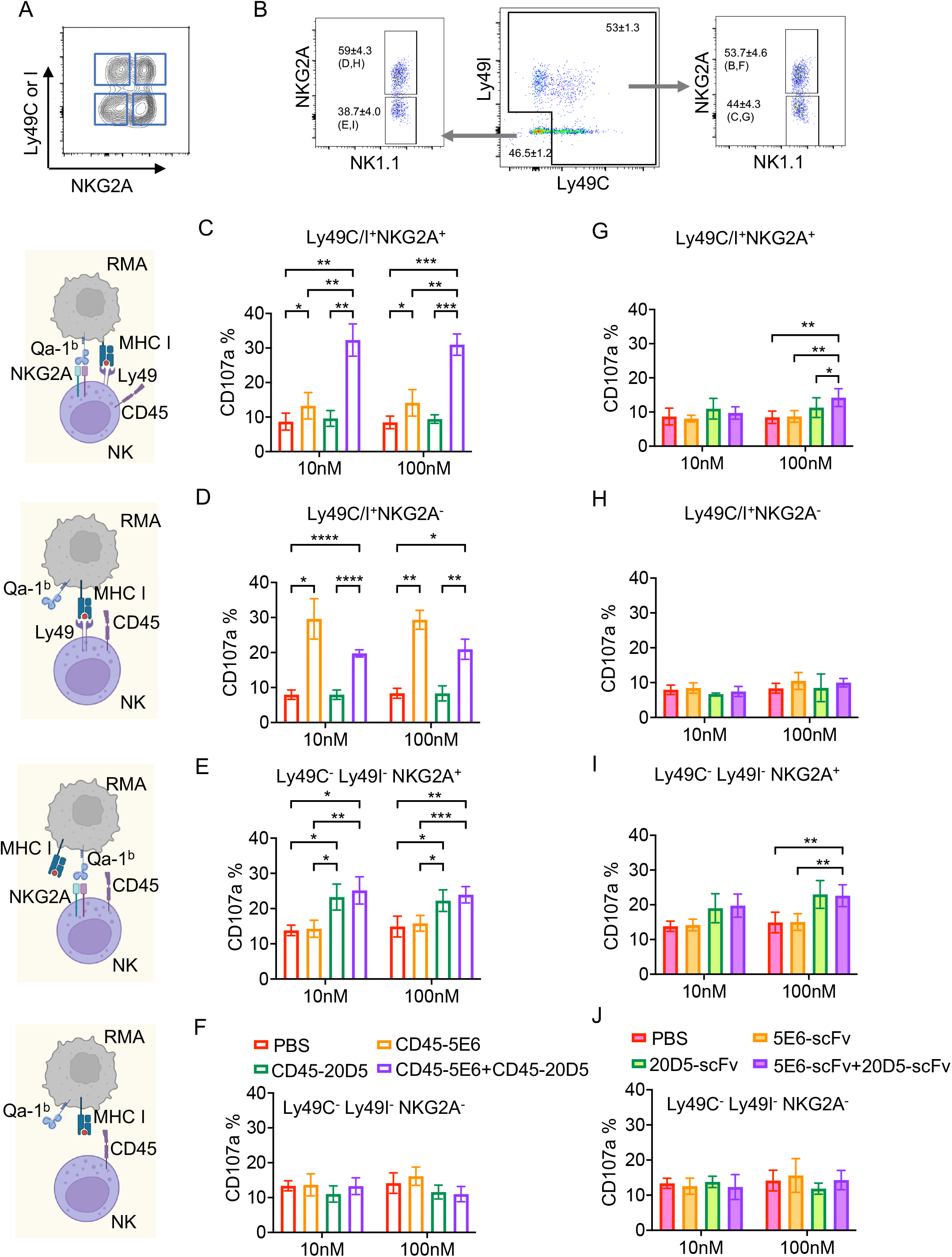
Combination of CD45-5E6 and CD45-20D5 potentiates NK activation. (A) FACS distributions showing the NKG2A, Ly49C or Ly49I expression in resting splenic NK cells. (B) Four different NK cell subpopulations defined by expression and co-expression of Ly49I, Ly49C and NKG2A in WT C57BL/6 mice that were pretreated with 200 μg poly(I:C) for 2 days before stimulating the cells with RMA target cells in the presence or absence of CD45-NKRs or NKR-scFvs. RMA cells were precultured with 20 ng/ml IFNγ for upregulating Qa-1^b^ and were labeled with 1μM CTV. 1 × 10^6^ Splenocytes from poly(I:C) treated mice were cocultured with 2 × 10^5^ RMA Qa-1^b+^MHC I^+^ target cells in the presence of anti-CD107a-PE, brefeldin A, monensin, 5E6-scFv, CD45-5E6, 20D5-scFv, CD45-20D5, a combination of 5E6-scFv and 20D5-scFv, or a combination of CD45-5E6 and CD45-20D5 for 5h. The center plot shows costaining of gated NK cells for Ly49C and Ly49I. Gated NK cell populations that were positive for Ly49I and/or Ly49C, or negative for both, were then further gated based on NKG2A staining (right and left plots). (C-J) Each of the four populations examined was analyzed to determine the percentage of CD107a^+^ cells among Ly49C^+^ and/or Ly49I^+^ NK cells that also expressed NKG2A (Ly49C/I^+^NKG2A^+^) (C, G), Ly49C^+^ and/or Ly49I^+^ NK cells that did not express NKG2A (D, H), NKG2A^+^ NK cells that did not express Ly49C or Ly49I (E, I), and NK cells that did not express any of the three receptors (F, J). Bar graphs show mean ± SD (n=4) and were analyzed by two-way ANOVA. Multiple comparisons were corrected using Tukey test. Data are representative of three independent experiments. *P < 0.05; **P < 0.01; ***P < 0.001; ****P < 0.0001.

Not surprisingly, NK cells that lacked all three receptors failed to respond to any of these molecules, in CD45-NKR or scFv format (Fig 6F, J, Fig. S6H). Furthermore, the responses of the various subpopulations of NK cells to MHC I-deficient target cells was not much affected by the CD45-NKRs, alone or in combination (Fig. S7), consistent with the analysis of NK cell killing, which showed that the effects of CD45-NKRs were much weaker when MHC I deficient target cells were employed.

The synergistic effect of the CD45-NKR combinations on some NK cells suggests that combining CD45-NKRs that target these receptors is a potential strategy to improve their immunotherapeutic efficacy. Not only does the combination activate a subpopulation of NK cells that neither CD45-NKR alone activates well (the approximately 30% of NK cells that co-expressed NKG2A and Ly49C/I), but each CD45-NKR activates unique NK subsets, all of which can be simultaneously activated by the CD45-NKR combination. As a result of these two features, the subpopulations that respond to the CD45-NKR combination together comprise approximately 80% of NK cells, whereas the populations that respond to each CD45-NKR separately comprise only 21-32% of NK cells.

## Discussion

Induced proximity of cell surface proteins is gaining increasing interest for modulating cell surface receptor signaling. Strategies for down-tuning receptor signaling include recruitment of cell surface E3 Ub ligases, cell surface phosphatases, and mannose-6-phosphatase receptors(*39*, *44*, *45*). While E3 Ub ligase and MAN-6-P recruitment result in degradation of the target proteins, phosphatase recruitment exploits the intrinsically high activity of phosphatases such as CD45 to intracellularly dephosphorylate targets within close proximity. On immune cells CD45 is the principal cell surface phosphatase and serves as a counterbalance to Src kinase actions on ITAM/ITIM/ITSM receptors, to control the phosphorylation state of many important antigen receptors and checkpoint receptors (*46*, *47*). CD45 also plays other important roles in modulating TCR signaling by being segregated from the immunological synapse, thus potentiating the actions of Src kinases that act on the antigen receptor complex (*37*, *38*). Thus, ligation of inhibitory immunoreceptors to CD45 can result in not only reduction of their phosphorylation state directly, but also potentially enhance signaling through co-segregation with CD45 from the synapse, although this has not been directly shown. Additional mechanisms may also be at play.

Here we find that synthetic ligands that induced proximity between inhibitory mouse NKR and CD45 more effectively reverses inhibitory signaling by NKR, enhancing signaling and target cell killing by NK cells. We also show that the CD45-targeted NKG2A antagonist, CD45-20D5, potentiates CD8^+^ effector T cells in addition to NK cells. Upon binding to ligands such as MHC I or Qa-1^b^, inhibitory NK receptors negatively regulate NK activation signals. This study shows that CD45-NKR molecules enhance NK activity mainly by augmenting NK activation signals, including Erk S6 phosphorylation and upregulating cytokine receptor CD25. These data suggest that CD45-NKR efficacy reflects better antagonism of inhibitory signaling that prevents these activation signals in NK cells. The outcome manifests as improved target cell killing, degranulation, CD69 surface expression and secretion of effector cytokines.

The efficacy of the CD45-NKR molecules could reflect any of several non-mutually exclusive mechanisms, including more efficient termination of inhibitory signaling due to the recruitment of the phosphatase activity to the ITIM motifs of the NKR, an avidity effect of coligation to CD45, segregation of the NKR from the NK cell synapse with the target cells, or interference with tonic inhibitory signals from the NKR that do not depend on MHC I engagement.

Unlike PD-1, where clear evidence of tonic signaling exists (*39*), tonic signaling by inhibitory NKR has not been demonstrated (*48*). In our hands, CD45-NKR increased NK responses partially in some assays even when the target cells were MHC I-deficient. This finding is consistent with reduced tonic signaling, but also consistent with other mechanisms, such as inhibition due to engagement of MHC I on the NK cells themselves or on neighboring NK cells or other cells.

An interesting and potentially useful finding of our work is that separate inhibition of Ly49 or NKG2A with CD45-NKR largely failed to augment NK cell activation in the case of a large (~30%) subpopulation of NK cells that co-express NKG2A and Ly49C/I. These results are best explained by suggesting that these NK cells are fully inhibited by engagement of either NKG2A or Ly49C/I by Qa-1^b^ or MHC I, respectively, such that interfering with each separately is insufficient to boost NK cell activity. Consistent with this interpretation, combining the two CD45-NKRs activated this NK cell subpopulation synergistically.

CD45-5E6 molecules were effective with NK cells that expressed Ly49C/I but not NKG2A. Conversely, CD45-20D5 molecules were most effective with NK cells that expressed NKG2A but not Ly49C or Ly49I. Each of these populations represents only 25-30% of NK cells. Hence, targeting each of these receptor types separately may be insufficient to stimulate the most impactful NK cell responses. Combining the two CD45-NKRs led to activation of both of these subpopulations as well as the aforementioned subpopulation that co-expressed Ly49C/I and NKG2A. In total, these subpopulations represent approximately 80% of all NK cells. In contrast, the subpopulations that respond to CD45-20D5 or CD45-5E6 separately each represent only 21%-32% of NK cells. These considerations lead to the proposal that combining CD45-NKR that interfere with both types of receptors may provide much stronger efficacy than either alone in cancer immunotherapy.

The results of our studies on mouse CD45-NKR fusion proteins suggest that evaluation of this strategy in the context of human NKR antagonists may be of benefit in cancer immunotherapy. Two anti-KIR blocking antibodies, Lirilumab and IPH4102 have been tested in clinical trials; however, Lirilumab has not shown good efficacy in the treatment of acute myeloid leukaemia (AML) (*33*, *34*). Our findings suggest that the efficacy of Lirilumab in cancer therapy might be enhanced by CD45 ligation, although this remains to be tested. Collectively, this work in the mouse system uncovers new strategies to design NKR antagonists that warrant testing in human systems.

## Methods

### Cells

Primary NK cells, primary T cells, RMA, K562, 721.221 cell lines were kept in a humidified incubator at 37°C with 5% CO2 and cultured in complete RPMI1640 medium supplemented with 2 mM GlutaMAX (Invitrogen), 10% FBS (Sigma),10 mM HEPES pH 8.0 (Thermo Fisher), 1 mM sodium pyruvate (Gibco) and 50U/ml penicillin and streptomycin (Thermo Fisher). B16F10 and B16-OVA cells were cultured in complete DMEM medium containing 10% FBS, 50U/ml penicillin and streptomycin, 2 mM GlutaMAX, and dissociated with TrypLE™ Express Enzyme (Thermo Fisher) when splitting every other day. Expi293 cells (female-derived kidney cell line) were grown in Expi293™ Expression Medium (Thermo Fisher) at 37°C, 125 rpm, 5% CO2 atmosphere with 80% humidity in flasks with ventilated caps. Hi5 cells were cultured in ESF 921 media with 10 mg/L of gentamicin sulfate at 27 °C on a shaker. SF9 cells were grown in SF900 media, which contain 10% FBS, 10 mg/L of gentamicin sulfate and 2 mM GlutaMAX at 27 °C.

### Mice

C57BL/6 wild-type mice and C57BL/6-Tg (TcraTcrb) 1100Mjb/J (OT-I) transgenic mice were purchased from the Jackson Laboratory. All mice were housed in animal facility at Stanford University and maintained according to protocols approved by Stanford University Institutional Animal Care and Use Committee on Animal Care.

### Design of CD45-targeted NK receptor antagonists

The hybridomas of clone 5E6 was kindly gifted from Dr. Wayne M. Yokoyama, clone 20D5 and 16A11 were kindly gifted from Dr. David Raulet. The VL and VH sequences of 5E6, 20D5 and 16A11 antibodies were extracted from the sequencing results provided by GenScript PROBIO. A mouse CD45-NKR molecule is composed of an anti-mouse CD45 nanobody(*49*) fused to an anti-mouse NK receptor scFv using a GGGGTGGS linker.

### Protein expression and purification from insect cells

SF9 cells were transfected with plasmids making P0 viruses. P1 viruses were made from P0 infected SF9 cells. 1 or 2L of Hi5 cells at 2 × 10^6^/ml were infected with 1-3 ml P1 viruses. After 2-3 days of protein production, the supernatants were harvested and purified by incubating with Ni-NTA resin for 4h at room temperature. The resin was collected and washed with 1 x HBS containing 20mM imidazole. Proteins were eluted with 1 x HBS containing 250mM imidazole, filtered and purified in size-exclusion chromatography (SEC). Protein fractions were collected and concentrated. After snap-freezing in liquid nitrogen, the proteins were stored in a −80°C freezer. The purity and size of the proteins were further confirmed by SDS-PAGE and Coomassie Blue staining.

### Generation of mouse Effector NK cells

Spleens were dissected and minced from C57BL/6 mice. Red blood cells (RBC) were lysed with 1 x ACK lysis buffer (BioLegend). The NK cells were purified with NK cell isolation kit (Miltenyi Biotec) according to manufacturer’s instructions, cultured in complete RPMI1640 medium with 100IU/ml murine IL-2 at 0.2-0.5 × 10^6^/ml. Fresh medium was added every other day. The cells were harvested on day 5 or 6, stained with flow antibodies, YLI-90 for anti-Ly49I, 16A11 for anti-NKG2A, and sorted for live CD3^-^CD49b^+^Ly49^+^ or NKG2A^+^ NK cells using a SH800S cell sorter (Sony). The sorted cells were allowed to recover for 2 days before using as effector cells in the co-culture assays.

### Antibodies for Flow Cytometry

Mouse experiments: Antibodies to CD16/32 (93), CD107a (1D4B, APC-Cy7), CD69 (H1.2F3, BV785), GM-CSF (MP1-22E9, PE-Cy7), CD49b (DX5, PE) and CD94 (18d3, PerCP/Cy5.5) were purchased from BioLegend. Anti-Ly49I (YLI-90, FITC), anti-CD3 (145-2C11, PerCP/Cy5.5), anti-CD8 (53-6.7, PE-Cy7), anti-MHC I (AF6-88.5.5.3, APC) and Goat anti-Mouse IgG Cross-Adsorbed Secondary Antibody, Alexa Fluor 647 were obtained from Thermo Fisher. Antibodies to Ly49C and Ly49I (5E6, FITC), Qa-1^b^ (6A8.6F10.1A6, PE), IFNγ (XMG1.2, Alexa Fluro 647) were purchased from BD Bioscience. Anti-HA (6E2, Pacific Blue) was purchased from Cell Signaling Technology. Anti NKG2A (705829, Alexa Fluro 647) was purchased from R&D Systems. The 4LO3311 hybridoma (anti-Ly49C) was a generous gift from S. Lemieux (Quebec, Canada), and the mAb was purified from CELLine classic 1000 (Integra Biosciences, Chur, Switzerland) supernatant, then conjugated to biotin according to standard methods. Biotin-conjugated mAb were detected with streptavidin PE-Cy7 (BioLegend).

### Antibodies for Cell Activation

Anti-mouse NKG2A (16A11, purified) was purchased from BioLegend. Anti-mouse NKG2A (20D5, purified) and anti-mouse CD28 (37.51, purified) were purchased from BioXcell. Anti-mouse Ly-49C and Ly-49I (5E6, purified) was purchased from BD Bioscience.

### Flow cytometry

Cells were washed in FACS buffer (2% FBS in PBS) and incubated with Fc receptor blocking antibody for 10 min, followed by staining with anti CD3, NK1.1, Ly49I, NKG2A for 20min at 4°C. Intracellular staining of IFNγ, CCL5 or IL-2 was performed according to the protocol of eBioscience™ Intracellular Fixation & Permeabilization Buffer Set (Cat.88-8824-00, Thermo Fisher). Briefly, cells were fixed in fixation buffer for 20 min at room temperature. After washing with FACS buffer, cells were resuspended, washed in 1X permeabilization buffer and stained with anti IFNγ, CCL5 or IL-2 antibodies for 20min at 4°C. Cells were resuspended in FACS buffer after washing. Flow cytometry was performed using a CytoFlex (Beckman Coulter). Data were analyzed using FlowJo software.

### NK cytotoxicity assay

For CD45-20D5, RMA cells were preincubated with 20 ng/ml IFNγ for 2 days. For CD45-5E6, RMA cells did not require IFNγ stimulation. Target cells were labeled with 1 μM cell trace violet (CTV) (Thermo Fisher) for 20 min at 37°C. Effector NK cells were co-cultured with 10,000 target cells at different effector/target ratios in the presence of 100nM CD45-NKR molecules. After 4 hours, the cells were spun down, washed with PBS, and assayed with Annexin V/7-AAD staining kit (BioLegend). The cells were stained with Annexin V diluted in 1x Annexin V buffer for 15 minutes and incubated with 2x 7-AAD for 5 minutes. The cells were analyzed in a CytoFlex flow Cytometer (Beckman Coulter).

### NK degranulation and activation assays

Effector NK cells at 1 × 10^6^/ml were co-cultured with target cells at 1:2 ratio for 4h, anti-CD107a (1D4B, APC-Cy7), brefeldin A (BD), monensin (BD) and serial dilutions (6x) of CD45-NKR and antibody controls were added to the coculture wells at the beginning of the assay. The cells were stained with a panel of antibodies for gating NK subsets and testing NK activation. Production of effector cytokines was tested by intracellular staining of IFNγ. The cells were analyzed in a Cytoflex Flow Cytometer. FACS results were plotted by FlowJo (TreeStar) and Prism 9.3.1 (GraphPad).

### Generation of mouse Effector OT-I cells

Splenocytes were harvested from OT-I mice. CD8^+^OT-I cells were purified by CD8^+^ T Cell Isolation kit (Mitenyi Biotec) and cultured for 7 days in complete RPMI1640 supplemented with 0.5ug/ml anti CD28 (37.51, purified), 1ug/ml OVA257-264 peptide (GenScript) and 100IU/ml mIL-2. The cells were splitted when the concentration reach 1.5 × 10^6^/ml. NKG2A expressing level was confirmed by FACS staining.

### OT-I degranulation and activation assays

B16-OVA cells were preincubated with 20ng/ml IFNγ for upregulation of MHC I and Qa-1^b^. Target cells were labeled with 1μM CTV prior to coculture. In the presence of anti-mouse CD107a, brefeldin A (BD) and monensin (BD), effector OT-I cells at 1 × 10^6^/ml were co-cultured with target cells at a ratio of 1:2 by adding serial dilutions (6x) of CD45-20D5 and 20D5-scFv to the coculture wells. 4h later, the cells were stained with anti-CD3, CD8, NK1.1, NKG2A, CD69, CD25. Anti-NKG2A (705829, Alexa Flour 647) was used for gating NKG2A^+^ OT-I cells. Effector cytokine production was tested by intracellular staining for IFNγ and IL-2.

### NK responsiveness assay

At day 0, C57BL/6 mice were treated with 200ug poly(I:C) (Invivogen) for 2 days. RMA cells were treated with 20ng/ml IFNγ for upregulating Qa-1^b^. At day 2, target cells were labeled with 1μM cell trace violet (CTV) (Thermo Fisher) for 20min at 37°C. The splenocytes were harvested. Splenic cells at 1 × 10^6^ were co-cultured with target cells at 1:2 effector to target ratio in the presence of anti-CD107a (Biolegend), brefeldin A and monensin and a serial dilutions (6x) of CD45-NKR molecules for 5 hours. were added at the beginning of the assay. NK subpopulations were stained and gated by using a panel of antibodies prior to detection. Anti-Ly49I (YLI-90, FITC) and anti-NKG2A (705829, Alexa Flour 647) were used for gating Ly49I^+^ and NKG2A^+^ cells. Cells were analyzed in a Cytoflex Flow Cytometer. NK responsiveness was assayed by cell surface staining of CD107a and intracellular staining IFNγ. Results were plotted by FlowJo and Prism 9.3.1.

### Combination treatment assay

RMA cells were treated with 20ng/ml IFNγ to induce upregulation of Qa-1^b^. 1 × 10^6^ Splenocytes from poly(I:C) treated mice were harvested and co-cultured with 2 × 10^5^ RMA or RMA-*B2m^-/-^* cells in the absence or presence of 10nM or 100nM 5E6-scFv, CD45-5E6, 20D5-scFv, CD45-20D5, 5E6-scFv+20D5-scFv, CD45-5E6+CD45-20D5 for 5 hours. Anti-CD107a, brefeldin A and monensin were added to the coculture wells before the experiment. The cells were stained with FACS antibodies and analyzed in Cytoflex. Anti-Ly49I (YLI-90, FITC), anti-Ly49C (4LO3311, biotin-conjugated antibody followed by secondary staining with Streptavidin-PE-Cy7) and anti-NKG2A (705829, Alexa Flour 647) were used for gating Ly49I^+^, Ly49C^+^ and NKG2A^+^ cells. NK responsiveness in Ly49C^+^Ly49I^+^NKG2A^+^, Ly49C^+^Ly49I^+^NKG2A^-^, Ly49C^+^Ly49I^-^NKG2A^+^, Ly49C^-^Ly49I^+^NKG2A^+^, Ly49C^+^Ly49I^-^NKG2A^-^, Ly49C^-^Ly49I^-^NKG2A^-^, Ly49C^-^Ly49I^+^NKG2A^+^ and Ly49C^-^Ly49I^-^NKG2A^-^ cells was tested by cell surface staining of CD107a. Results were plotted by FlowJo and Prism 9.3.1.

### Statistical analysis

Statistics were performed using Prism v9.3.1(350) (GraphPad Software). Data are expressed as mean ± standard deviation unless otherwise indicated. We used students t-test to analyze experiments with two groups, one-way ANOVA followed by Dunnett’s multiple comparisons tests and two-way ANOVA followed by Tukey’s multiple comparisons tests to analyze experiments with more than two groups. Significance is indicated as follows: *p < 0.05; **p < 0.01; ***p < 0.001, ****p<0.0001.

## Acknowledgements

We thank Thorbald van Hall for WT RMA and RMA-Qa-1^b^ KO cells. We acknowledge Dario Campana for sharing the K562-GpHLA-E cells. This work was supported by 2 R01 CA177684 06A1, and Emerson Collective to KCG, and by NIH grant R01 AI113041 to DHR. Portions of the schematics in the figures were generated by BioRender.

## Data availability

Data that support the findings of this study are available from the corresponding author upon reasonable request. Source data are provided with this paper.

---

**Figure S1.**
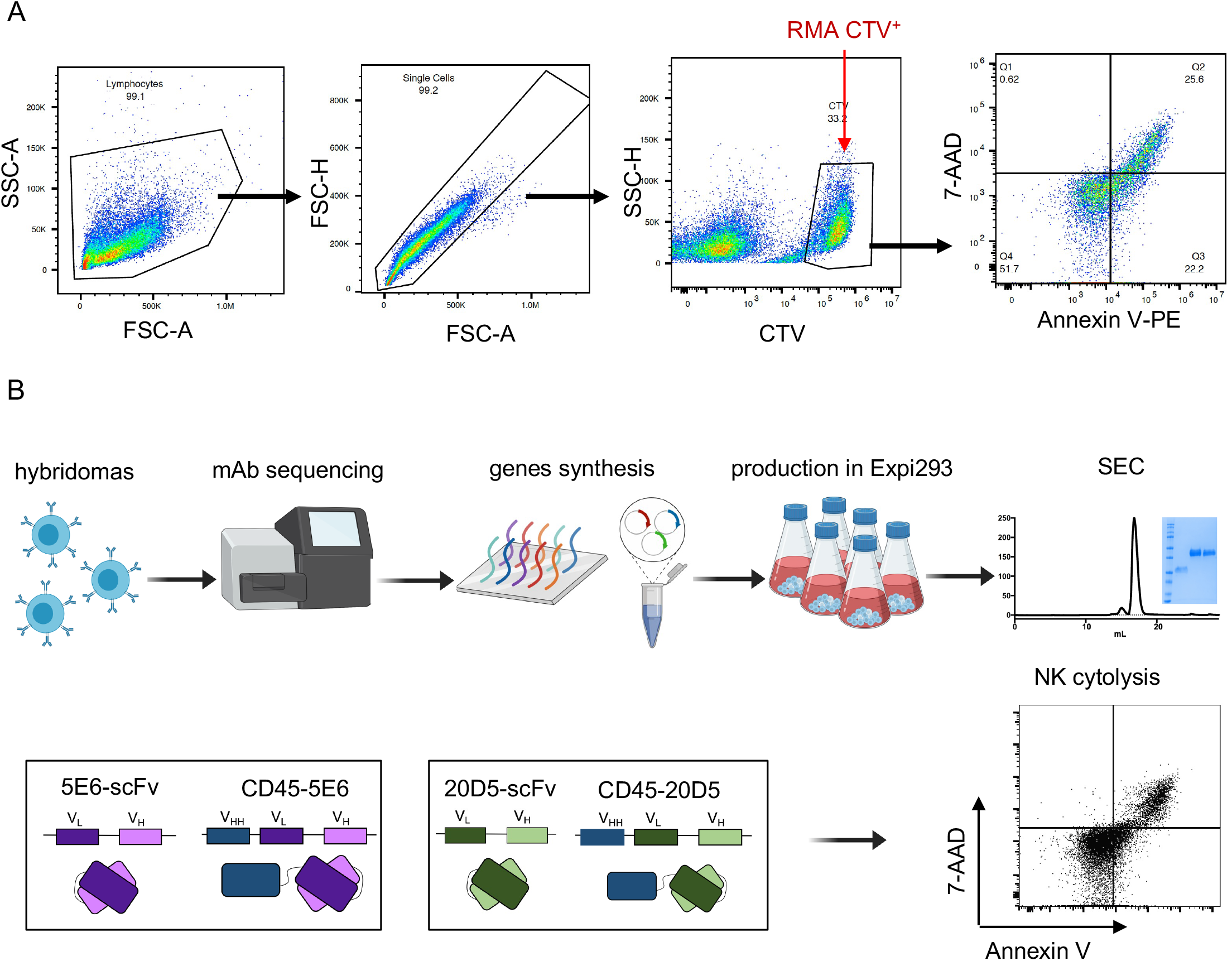
Engineering strategy of inhibitory CD45-NKR. (A) Gating strategy of NK cytolysis by annexin V/7-AAD staining. (B) Schematic pipeline of re-engineering of NKR antibodies, protein expression and NK-cell functionality testing.

**Figure S2.**
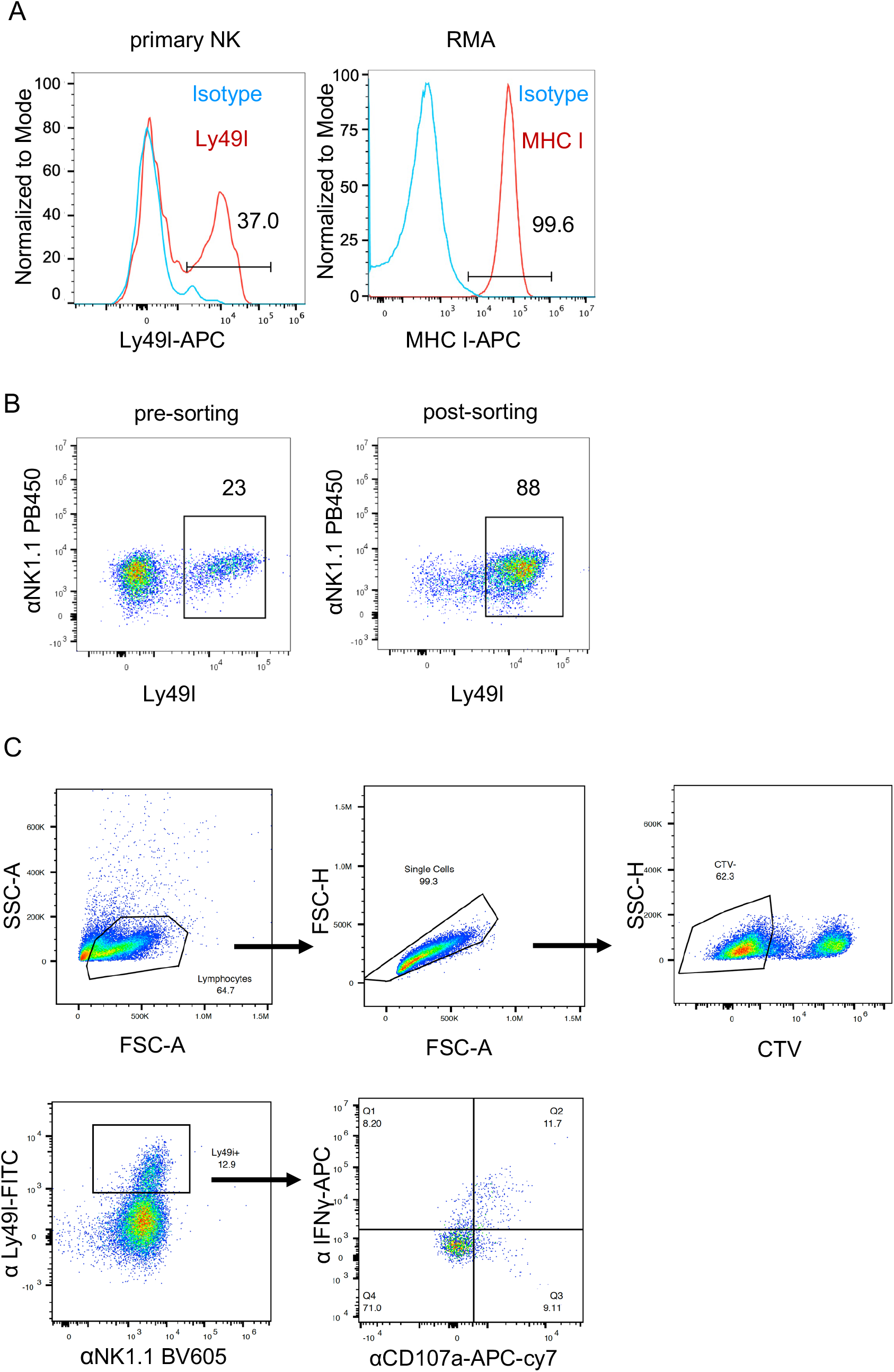
Validating surface expression of both Ly49I and MHC I. (A) The resting NK cells were stained with anti-Ly49I (YLI-90, FITC). The RMA cells were stained with anti-MHC I (AF6-88.5.5.3, APC). Histogram showing surface expression of Ly49I in NK and MHC I in RMA cells. (B) IL-2 expanded NK cells were stained with anti-Ly49I (YLI-90, FITC). Examples showing surfacing expression of Ly49I in before and after Ly49I sorting. (C) Gating scheme for NK degranulation and activation. Ly49^+^ NK cells were stained and gated with anti Ly49I (YLI-90, FITC).

**Figure S3.**
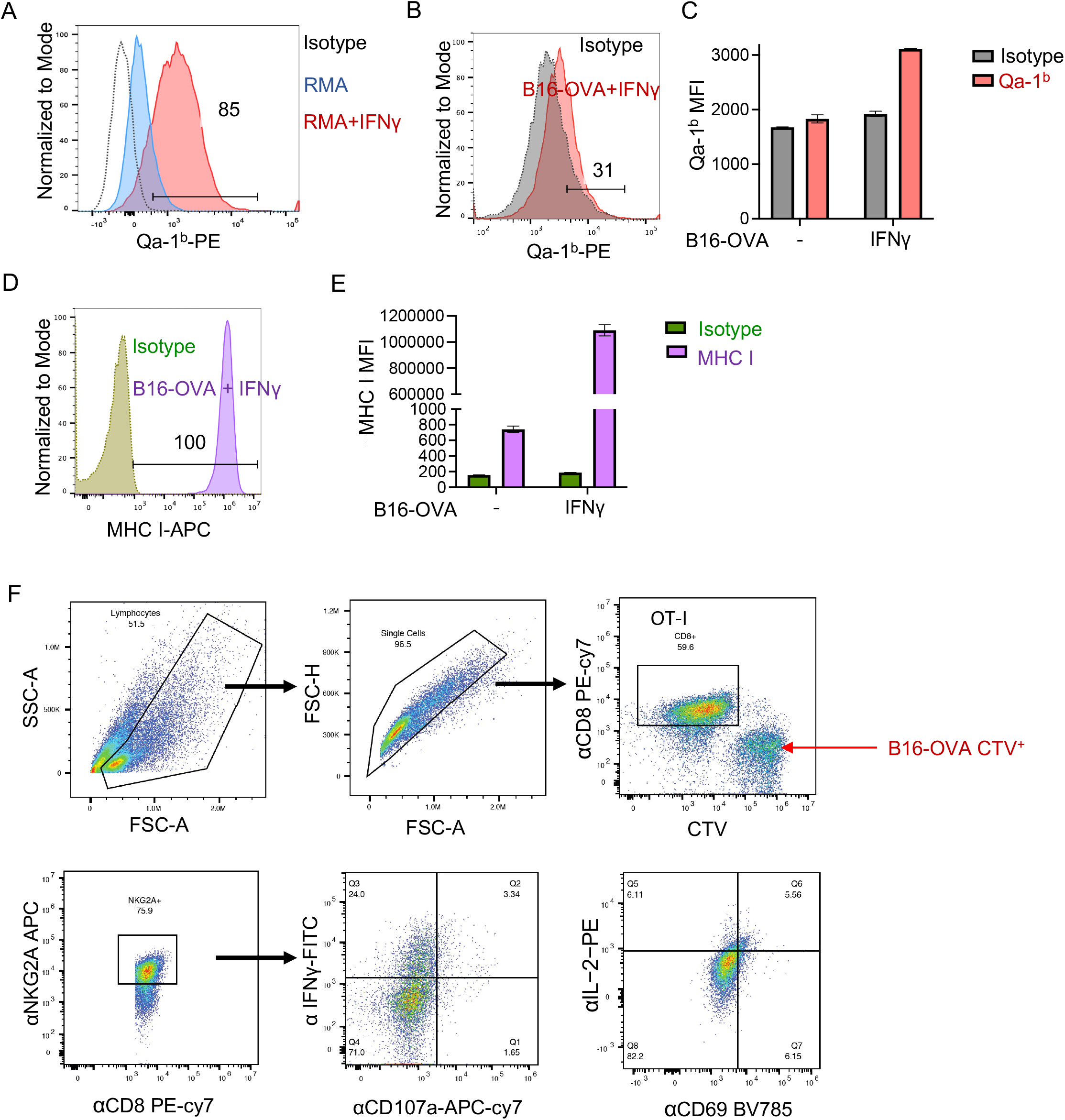
Validating surface expression of both Qa-1^b^ MHC I in target cells. (A) Histogram showing surface expression of Qa-1^b^ in IFNγ treated RMA cells. (B) Histogram showing surface expression of Qa-1^b^ in IFNγ treated B16-OVA cells. (C) Geometric mean fluorescence intensity (MFI) of Qa-1^b^ in IFNγ treated B16-OVA cells. (D) Histogram showing surface expression of MHC I in IFNγ treated B16-OVA cells. (E) Geometric mean fluorescence intensity (MFI) of MHC I in IFNγ treated B16-OVA cells. (F) Gating scheme for OT-I degranulation and activation.

**Figure S4.**
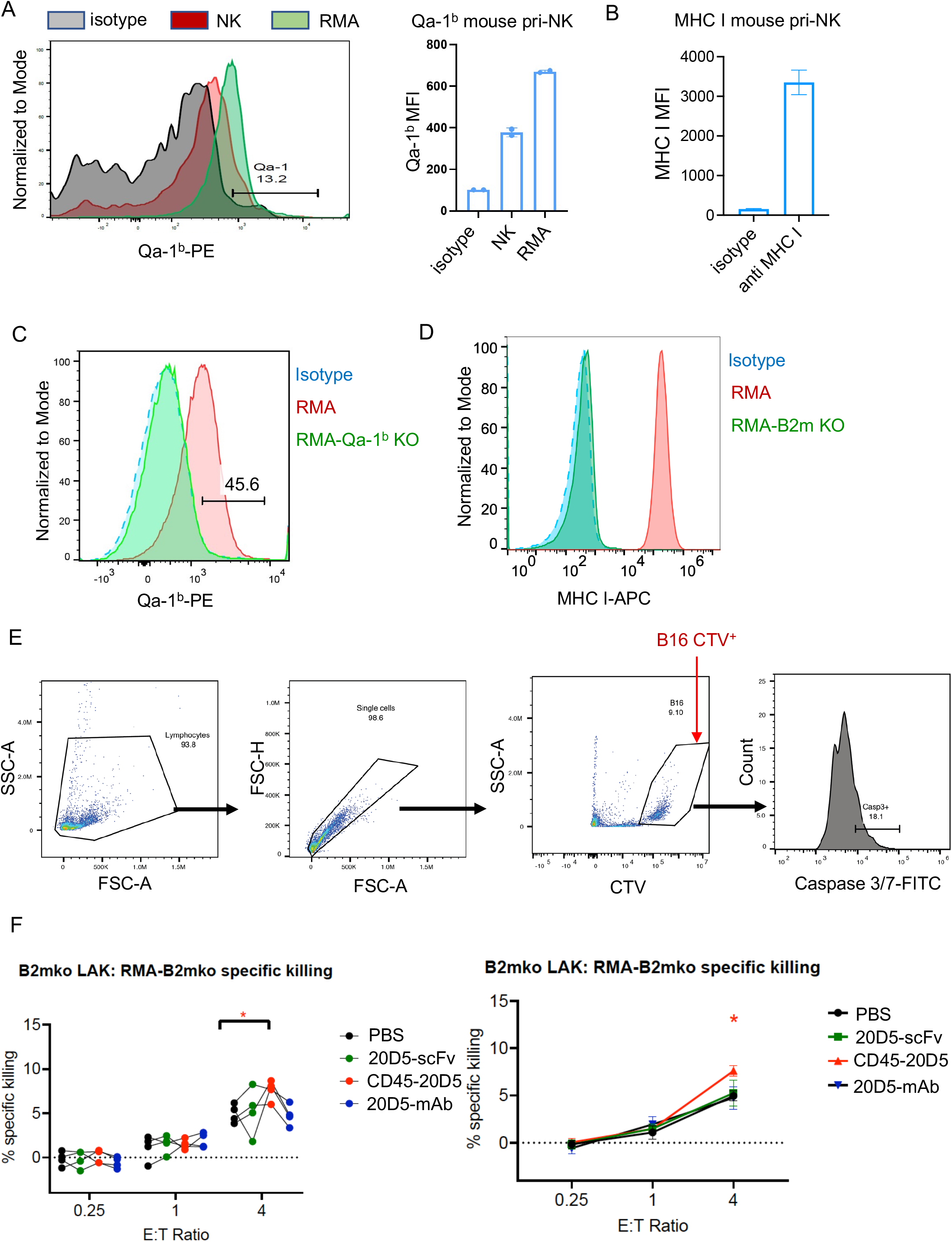
Validating surface expression of both Qa-1^b^ MHC I in RMA-Qa-1^b^ KO and RMA-B2m KO cells. (A) Histogram showing surface expression of Qa-1^b^ in RMA-Qa-1^b^ KO and IFNγ treated RMA cells. (B) Histogram showing surface expression of MHC I in RMA-B2m KO and RMA cells. (C) Gating scheme for NK cytotoxicity of B16 cells by detecting capase-3 activity. (D) Cytolysis of RMA cells by B2m KO NK cells In the presence or in the absence of 20D5-scFv and CD45-20D5 at 100nM. The Cytolysis were quantified by Annexin V/7-AAD assay.

**Figure S5.**
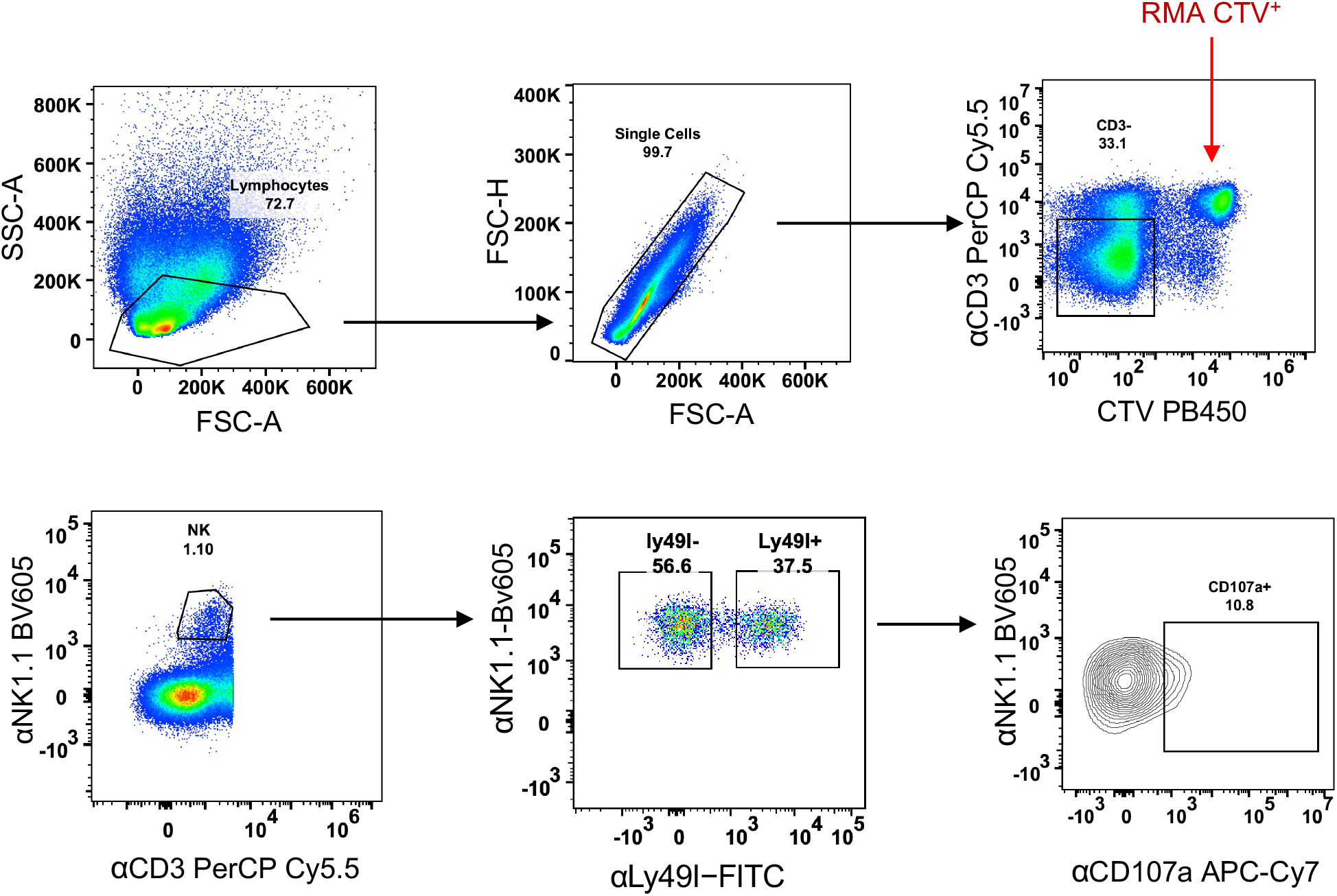
Cross talk between Ly49 and NKG2A receptors. Gating scheme for NK responsiveness by detecting CD107a. WT mice were treated with 200μg poly(I:C) for 2 days. RMA cells were labeled with 1μM CTV. Splenocytes from poly(I:C) treated mice were harvested and cocultured with target cells at 1:2 ratio in the presence of anti-CD107a-APC-Cy7, brefeldin A, monensin, NKR scFv and CD45-NKR for 5h. The cells were stained with a panel of surface staining antibodies, analyzed in CytoFlex flow cytometer.

**Figure S6.**
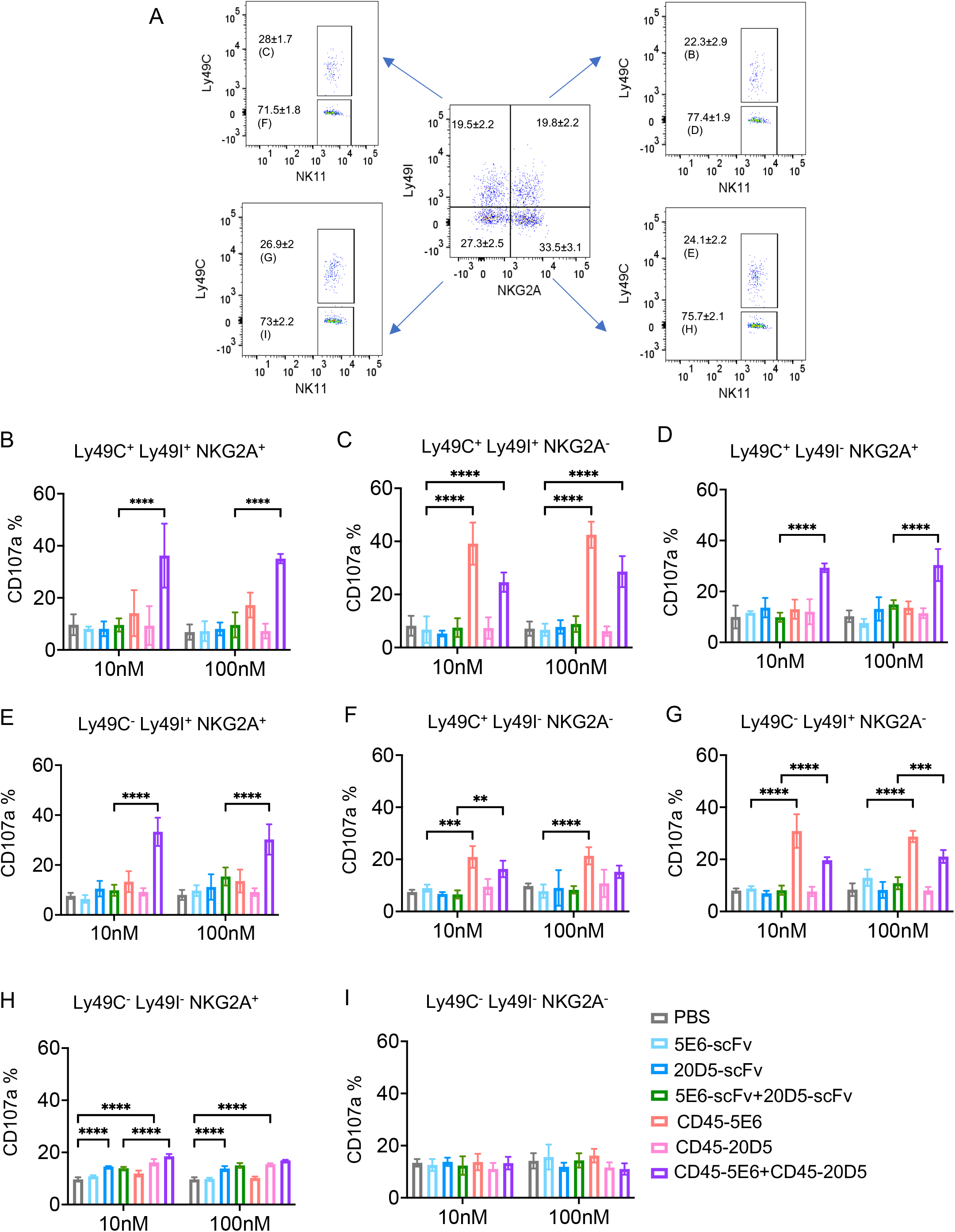
Combination effect of CD45-5E6 and CD45-20D5 on NK cells stimulated by target cells expressing MHC I and Qa-1^b^. (A) Eight different NK cell subpopulations defined by expression and co-expression of Ly49I, Ly49C and NKG2A in WT C57BL/6 mice that were pretreated with 200 μg poly(I:C) for 2 days before stimulating the cells with RMA target cells in the presence or absence of CD45-NKRs or scFvs. RMA cells were precultured with 20 ng/μl IFNγ for upregulating Qa-1^b^ and were labeled with 1μM CTV. 1 × 10^6^ Splenocytes from poly(I:C) treated mice were cocultured with 2 × 10^5^ RMA Qa-1^b+^MHC I^+^ target cells in the presence of anti-CD107a-PE, brefeldin A, monensin, 5E6-scFv, CD45-5E6, 20D5-scFv, CD45-20D5, a combination of 5E6-scFv and 20D5-scFv, or a combination of CD45-5E6 and CD45-20D5 for 5h. Representative flow cytometry plots for gating 8 different NK cell subsets expressing none, any one, any pairs or all of the Ly49C, Ly49I, and NKG2A receptors are shown. (B-I) Each of the eight populations examined was analyzed to determine the percentage of CD107a^+^ cells in Ly49C^+^Ly49I^+^NKG2A^+^ (B), Ly49C^+^Ly49I^+^NKG2A^-^ (C), Ly49C^+^Ly49I^-^NKG2A^+^ (D), Ly49C^-^Ly49I^+^NKG2A^+^ (E), Ly49C^+^Ly49I^-^NKG2A^-^ (F), Ly49C^-^Ly49I^+^NKG2A^-^ (G), Ly49C^-^Ly49I^-^NKG2A^+^(H) and Ly49C^-^Ly49I^-^NKG2A^-^ (I) NK cells. Bar graphs show mean ± SD and were analyzed by two-way ANOVA. Multiple comparisons were corrected using Tukey test. Data are representative of two independent experiments. *P < 0.05; **P < 0.01; ***P < 0.001; ****P < 0.0001.

**Figure S7.**
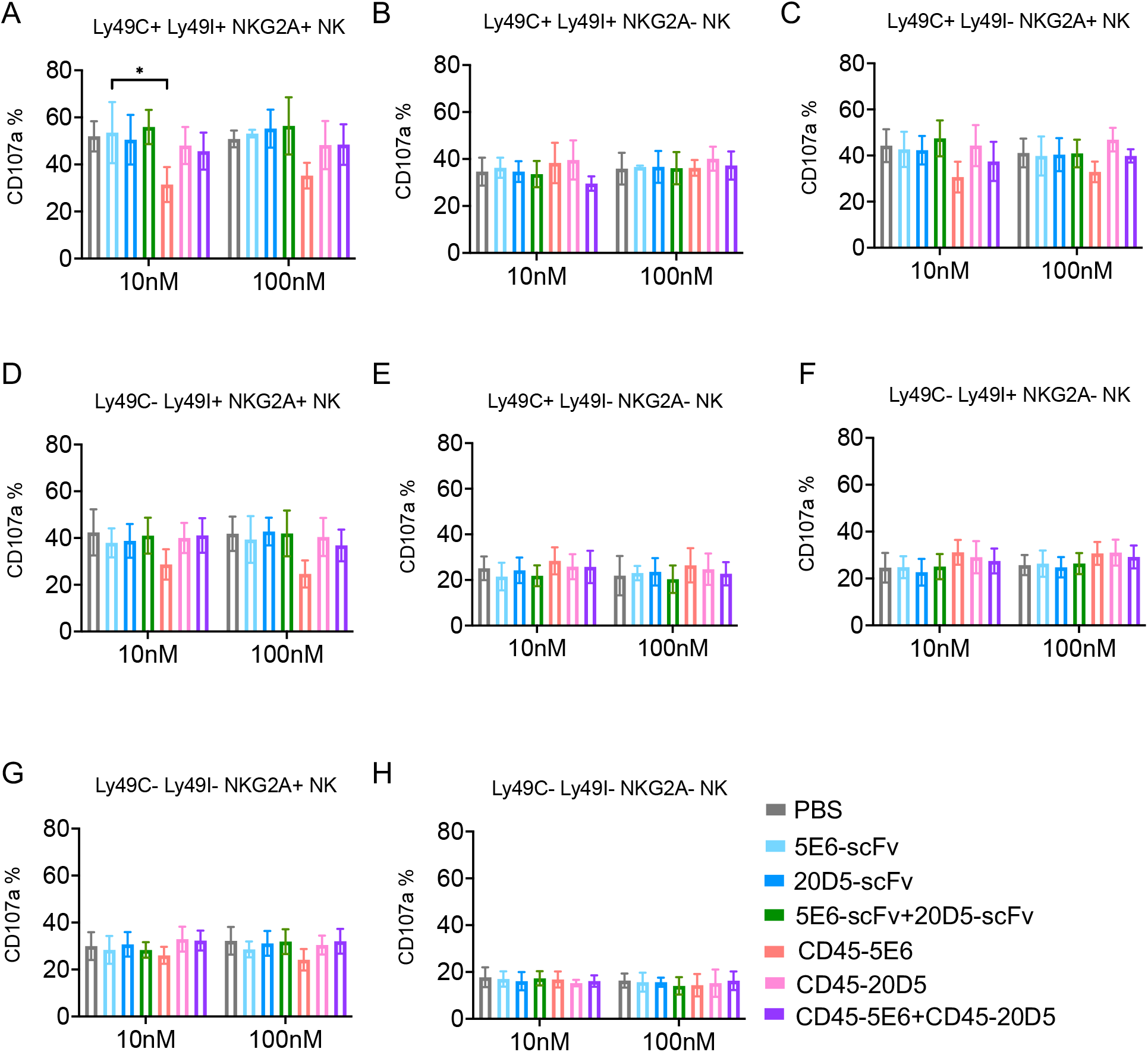
Combination effect of CD45-5E6 and CD45-20D5 on NK cells stimulated by MHC I-deficient target cells. (H) NK cells were stimulated with RMA-*B2m* KO tumor cells (which also lack Qa-1^b^) in the presence of 5E6-scFv, CD45-5E6, 20D5-scFv, CD45-20D5, a combination of 5E6-scFv and 20D5-scFv, a combination of CD45-5E6 and CD45-20D5. Quantification of NK cell responsiveness by Ly49C^+^Ly49I^+^NKG2A^+^ (A), Ly49C^+^Ly49I^+^NKG2A^-^(B), Ly49C^+^Ly49I^-^NKG2A^+^ (C), Ly49C^-^Ly49I^+^NKG2A^+^ (D), Ly49C^+^Ly49I^-^NKG2A^-^ (E), Ly49C^-^Ly49I^+^NKG2A^-^ (F), Ly49C^-^Ly49I^-^NKG2A^+^ (G) and Ly49C^-^Ly49I^-^NKG2A^-^ (H) NK cells. Bar graphs show mean ± SD and were analyzed by two-way ANOVA. Multiple comparisons were corrected using Tukey test. Data are representative of two independent experiments. *P < 0.05; **P < 0.01; ***P < 0.001; ****P < 0.0001.

